# Disequilibrium of fire-prone forests sets the stage for a rapid decline in conifer dominance during the 21^st^ century

**DOI:** 10.1101/163899

**Authors:** Josep M. Serra-Diaz, Charles Maxwell, Melissa S. Lucash, Robert M. Scheller, Danelle M. Laflower, Adam D. Miller, Alan J. Tepley, Howard E. Epstein, Kristina J. Anderson-Teixeira, Jonathan R Thompson

## Abstract

As trees are long-lived organisms, the impacts of climate change on forest communities may not be apparent on the time scale of years to decades. While lagged responses to environmental change are common in forested systems, potential for abrupt transitions under climate change may occur in environments where alternative vegetation states are influenced by disturbances, such as fire. The Klamath mountains (northern California and southwest Oregon, USA) are currently dominated by carbon rich and hyper-diverse temperate conifer forests, but climate change could disrupt the mechanisms promoting forest stability– regeneration and fire tolerance— via shifts in the fire regime in conjunction with lower fitness of conifers under a hotter climate. Understanding how this landscape will respond to near-term climate change (before 2100) is critical for predicting potential climate change feedbacks and to developing sound forest conservation and management plans. Using a landscape simulation model, we estimate that 1/3 of the Klamath could transition from conifer forest to shrub/hardwood chaparral, triggered by an enhanced fire activity coupled with lower post-fire conifer establishment. Such shifts were more prevalent under higher climate change forcing (RCP 8.5) but were also simulated under the climate of 1950-2000, reflecting the joint influences of early warming trends and historical forest legacies. Our results demonstrate that there is a large potential for loss of conifer forest dominance—and associated carbon stocks and biodiversity- in the Klamath before the end of the century, and that some losses would likely occur even without the influence of climate change. Thus, in the Klamath and other forested landscapes subject to similar feedback dynamics, major ecosystem shifts should be expected when climate change disrupts key stabilizing feedbacks that maintain the dominance of long-lived, slowly regenerating trees.

## INTRODUCTION

Climate change is expected to cause significant changes in forest carbon (C) cycling and species composition, with potentially profound effects given that forests globally harbor the majority of Earth’s terrestrial biodiversity and store ~45% of terrestrial organic C (Bonan, 2008; Pan *et al.*, 2011). How and when these shifts will manifest and how they will interact with land-use legacies and other mesoscale dynamics, is, however, not well understood. While statistical associations between species distributions and climate suggest the potential for rapid shifts in forests species ranges due to shifts in their associated suitable habitats (Iverson *et al.*, 2008; Serra-Diaz *et al.*, 2014), there is also mounting evidence of lagged responses (Bertrand *et al.*, 2011; Svenning & Sandel, 2013) that the literature has characterized as “debts” or borrowed time (Hughes *et al.*, 2013): climate change debt when species do not track their climatic environment fully (Bertrand *et al.* 2011), or resilience debt when species are not adapted to a given disturbance regime (Johnstone *et al.* 2016). Overall, these debts reflect that the current state of the forest communities may not readily reflect species climatic fitness, or equilibrium, but is rather a product of the past. Because of these lagged effects, predicting the actual responses of forests to near-term climate change (i.e. < 100 years) is thus an urgent priority for global change ecology.

Rapid changes to forest communities are likely when climate change induces shifts in disturbance regimes and extreme events, including changing patterns and occurrence of forest pests, the frequency and severity of drought mortality events (Allen *et al.*, 2010; Clark *et al.*, 2016), large-scale recruitment failures (Turner, 2010; Feddema *et al.*, 2013; Clark *et al.*, 2016; Johnstone *et al.*, 2016; Tepley *et al.*, 2017) or shifts in biotic interactions rendering shifted fitness of forest species in the community (Carnicer *et al.*, 2014). Alternatively, forest communities could be resilient, resistant, or simply experience gradual and delayed responses to climate change due to the long generation times of trees. These buffering mechanisms include demographic stabilization, CO_2_ fertilization, microclimatic buffering and landscape heterogeneity (Keenan *et al.*, 2011; Lloret *et al.*, 2012; De Frenne *et al.*, 2013; Bertrand *et al.*, 2016; Seidl *et al.*, 2016). Interactions between all these phenomena are complex and are an active focus of global change research (Franklin *et al.*, 2016).

In the view of forests as complex systems of multiple interacting mechanisms, rapid forest declines(<100 years) should be expected when key reinforcing feedbacks that maintain a stable community state are disrupted (Bowman *et al.*, 2015). For instance, both empirical and theoretical studies recognize that there is a range of climatic conditions in which tropical and subtropical forest and savanna may exist as alternative stable states modulated by vegetation-fire feedback dynamics (Staver & Levin, 2012). The persistence of these feedbacks is also a major concern in fire-prone temperate forests (Paritsis *et al.*, 2015; Johnstone *et al.*, 2016; Kitzberger *et al.*, 2016; Tepley *et al.*, 2016), and, under climate change, such feedbacks could either be expected to increase the frequency, size and/or severity of wildfires (Flannigan *et al.*, 2000; Westerling *et al.*, 2006),or reduce fire activity via negative feedbacks (Parks *et al.*, 2016; McKenzie & Littell, 2017). In addition, these new climate-change shifted fire regimes could interact with large-scale mortality with recruitment failures for some forest species (Enright *et al.*, 2015), and/or hamper potential re-colonization (Caughlin *et al.*, 2016; Tepley *et al.*, 2017). Therefore, a wide array of outcomes may be expected under climate change due to several mechanisms acting in concert.

Here we investigate the potential for rapid large-scale and fast transitions in forest communities due to interacting mechanisms due to climate change. We use the Klamath forest landscape (northern California and southern Oregon, USA) to portray potential shifts between high biomass conifer temperate forests (CON) and Mediterranean sclerophyll shrub, chaparral, and hardwood communities (SCH). The Klamath is a major carbon reservoir and an internationally recognized hotspot of botanical biodiversity (Briles *et al.*, 2005; Sawyer, 2007; Fig 1a). These vegetation types (conifer forests vs. hardwood-chaparral) are suggested to function at local scales as alternative stable states due to their self-stabilizing feedbacks involving biotic interactions and climate-fire (Petraitis & Latham, 1999; Odion *et al.*, 2010; Airey Lauvaux *et al.*, 2016) (Fig.1b). There are concerns about conifer regeneration failure in the Klamath, typically stemming from clear-cut logging (sensu Hobbs & Huenneke, 1992), and high severity wildfire, potentially exacerbated by climate change (Tepley *et al.*, 2017). The SCH state is comprised of highly pyrogenic SCH species (Brown & Smith, 2000) that inhibit conifer regeneration (Hobbs & Huenneke, 1992). Consequently, the SCH communities promote fire at a regime that hampers conifer regeneration (Thompson & Spies, 2010). Only when the fire-free interval is sufficiently long can conifers overtop the shrub layer and begin to dominate the site (Shatford *et al.*, 2007a). Indeed, mature conifer forests include (*Pseudotsuga menziesii*, *Calocedrus decurrens*, etc.) with fire-resistance adaptations that strengthen with age (e.g. bark thickening, crown base height, canopy shading; Agee, 1996; Shatford *et al.*, 2007b). At the mature conifer stage, a low severity fire regime is generally promoted, wherein surface fuels that affect only the understory and hamper ladder fuel development, reducing fire risk to the mature conifer overstory (Sensenig *et al.*, 2013).

**Figure 1:**
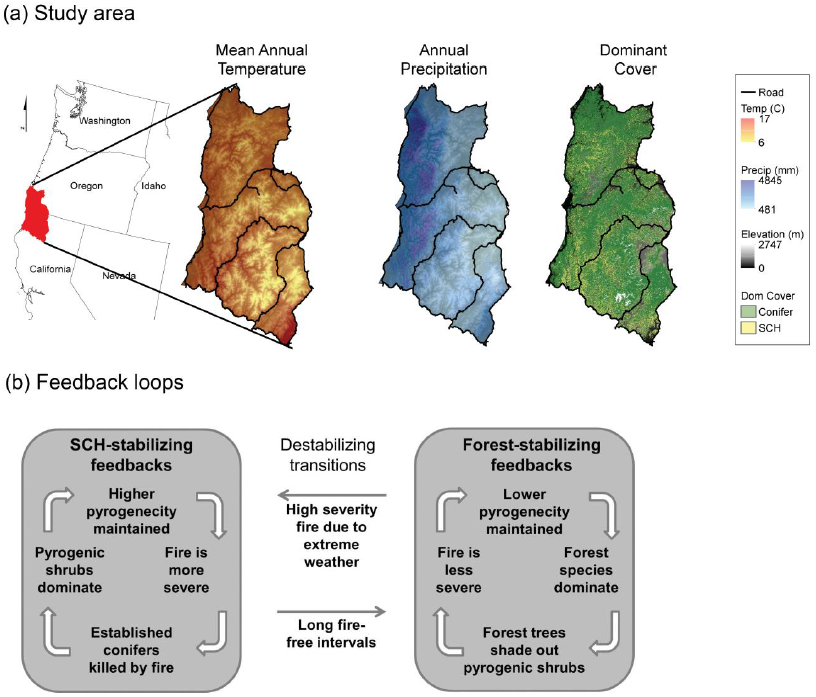
(a) Study area temperature, precipitation and initial forest dominant type (left to right), and (b) feedback loops that maintain the two forest community states (apapted from Odion et al 2010).

It is difficult to assess whether observations and local studies are only context dependent (e.g. post fire legacy effects (Johnstone *et al.*, 2016)), or if they could represent a major regional shift towards SCH. Because large-scale experimentation in forest is unrealistic, mechanistic models with explicit representation of species and adaptations to wildfire offer a tool to explore the individual and interacting effects of multiple climate change drivers as well as feedback mechanisms; and allow consideration of a breadth of spatial and temporal scales in forest community dynamics. The aim of this study is to test the hypothesis that a warming climate and increased fire activity will drive rapid conifer decline in the Klamath. We expected that, compared to forest dynamics driven by the current climate (baseline conditions), climate change would (1) increase fire activity (decrease fire rotation period), (2) slow conifer growth, and (3) trigger major declines of conifer dominance.

## METHODS

### Study area and species

The Klamath ecoregion is situated in the Pacific Northwest region of the United States (Figure 1) at the convergence of major North American floristic zones (Whittaker, 1960), and includes an exceptionally diverse flora, with strong components of sclerophyll broadleaf hardwood, coniferous, and herbaceous vegetation. Topography is mountainous and complex; elevation ranges from 100m to 2000 m asl. The climate is Mediterranean, with dry, warm summers and wet, mild winters. Mean January temperature is 6°C. Mean July temperature is 16°C. Mean annual precipitation is 2900 mm, with greater than 90% occurring in winter (Daly *et al.*, 2002, 2008). Before the onset of effective fire suppression (c. 1945) wildfire return intervals ranged from 6 to 60 years (Taylor & Skinner, 2003; Sensenig *et al.*, 2013), although the regionally the Klamath has supported various fire regimes and return intervals in the last 2000 years (Colombaroli & Gavin, 2010).

### Modeling framework

Our approach consists of modeling vegetation succession interactions with climate change to integrate both slow (e.g. forest stand development) and fast processes (e.g. disturbances) into projections of potential vegetation transitions at a regional scale (Walker & Wardle, 2014).

We simulated vegetation dynamics using the forest community model LANDIS-II v 6.1 (Scheller *et al.*, 2007); <u>http://www.landis-ii.org/</u>). This process-based model includes biophysical (climate and soils) and ecological processes (species interactions, dispersal) to simulate growth, mortality and regeneration at the species level. Different life history traits enable modeling inter- and intra- specific interactions (e.g. competition, facilitation) integrated with disturbance responses (e.g. resprouting). Species are simulated as age-cohorts that compete for and modify aboveground and belowground resources within each cell; disturbances and dispersal are spatially-explicit processes. The model has been widely used in both temperate forests and Mediterranean type ecosystems to investigate climate-fire-forest interactions (Syphard *et al.*, 2011; Loudermilk *et al.*, 2013; Liang *et al.*, 2016). We ran 9 model replications for each climate change and baseline climate scenarios, thus a total of 45 simulations. Simulation were run in a cluster for 25 days. We classified each cell as conifer state or shrubland-chaparral-hardwood state according to the most dominant species in each cell. Dominance classification was based on the functional identity (e.g. SCH or conifer) of the species with the highest biomass.

### Initial species distribution

Initial distribution of vegetation types was obtained from nearest neighbor imputation of forest inventory plots (Ohmann & Gregory, 2002; <u>https://lemma.forestry.oregonstate.edu/data</u>) and they were constant among replications and scenarios. This methodology assigns each cell within the study area a forest inventory and analysis plot (FIA; United States Forest Service, 2008) based on the similarity of environmental data and remote sensing image spectral properties. We pooled cells of initial resolution of 30 m in to cells of 270 m to align with the assumptions associated with modeling cohorts, though still retaining the capacity to identify species distributions (Franklin *et al.*, 2013). We chose to model the most prevalent target species as well as grouped shrubs species into functional groups according to their seed vs. resprouting behavior, as well as their N-fixation availability (see Succession section).

### Input geophysical data

Weather data was extracted from the United States Geological Service data portal (<u>http://cida.usgs.gov/gdp/</u>; June 2016). We used Maurer historical weather data for baseline conditions for the period 1949-2010 (Maurer *et al.*, 2002) and bias corrected constructed analogs v2 daily for climate change projections climate change projections. We chose four GCM - RCP that portray the breadth of climate change conditions modeled for the Klamath region—i.e. those spanning the widest range of average annual temperature and precipitation-predicted for the study area (ACCESS 8.5 (Ac85, hotter and drier); CNRM-CM5 8.5 (Cn85 hotter and wetter); CNRM-CM5 4.5 (slightly hotter and slightly wetter), MIROC5 2.6 (slightly hotter and slightly drier; Fig.S1).

Soil characteristics were obtained through the web soil survey (<u>http://websoilsurvey.sc.egov.usda.gov/App/HomePage.htm</u>) from the STATSGO2 database. We used the measures of soil organic matter content, and physical characteristics of soils that drive water balance dynamics and biogeochemistry in the model (e.g. drainage class, percentage of sand). Nitrogen deposition was obtained from the National Atmospheric Deposition Program database (<u>http://nadp.sws.uiuc.edu/</u>).

In order to harmonize different scales and data types among physical inputs we parsed the study area into ecoregions with similar environmental conditions (Serra *et al.*, 2011). We grouped different cells in environmental space according to their soil and climate characteristics through cluster analysis, using the *clara* function in the ‘cluster’ package (v 2.0.4) in R 3.3.3 (R library core team, 2017). We used soil drainage characteristics, field capacity, percentage of clay, percentage of sand, wilting point and soil organic content to define five soil regions. For the climate regions we defined 5 regions using precipitation, minimum temperature and maximum temperature. The final set of twenty-five ecoregions is derived from the combination of climate and soil regionalization sets (5×5).

### Succession

Vegetation succession is generated by competition for light and water within each cell, and is represented via C and N cycling through leaf, wood, fine and coarse roots by species and age cohorts using the LANDIS-II Century Extension (Scheller *et al.*, 2011). The succession model operates at a monthly scale and simulates growth as a function of water, temperature, and available nitrogen, while accounting for inter-cohort competition for light and space. Mortality is caused by disturbances (see next section), senescence (ongoing loss of trees and branches), and age (higher mortality rate when approaching species longevity). Regeneration and establishment are characterized by probabilities based on species-specific life history attributes that consider dispersal distances, sexual maturity, post-fire behavior (e.g. serotiny, resprouting), light, and water availability.

Tree and shrub species species-level parameters and functional type parameters. Functional groups were created by combining growth forms (e.g. hardwood, conifer, shrub), biogeochemical behavior (e.g. N-fixing) and seasonality in growth (e.g. evergreen, deciduous) (following (Loudermilk *et al.*, 2013) and (Creutzburg *et al.*, 2016a). Species and functional group parameters determine growth in response to climate and soil properties. We calibrated species’ cohort growth using two parameters: maximum monthly aboveground productivity, and large wood mass [g C/m2] at which half of the theoretical maximum leaf area is attained, using 950 Forest Inventory Analysis (FIA) plots representative of the different community types and climate gradients present in the study area. Full list of input parameters are available in Table S1-S9.

We calibrated the growth model to represent the biomass of the selected FIA plots. Accuracy assessment indicated that the model was able to capture species-specific biomass with an average deviation of 10% of the biomass in a plot for each species (Figure S3).

### Fire

Wildfire was simulated using the Dynamic Fires and Fuel extensions (Sturtevant *et al.*, 2009). This extension simulates the landscape-scale fire regime given input parameters of topography, fuel type, fuel condition and daily fire weather (see full model description and fuel implementation in Sturtevant *et al.*, 2009 and Syphard *et al.*, 2011). Weather changes the probability of ignition and fire spread rates according to fuel type, fuel condition and the probability of a sustained flame via fire weather indices (Beverly & Wotton, 2007). Fire causes mortality in tree cohorts according to their age and the difference between species’ fire tolerance and fire intensity. Higher fire intensity is required to kill older cohorts whereas younger cohorts are killed by less intense fire. Such fire intensity is related to the topography, the climate conditions and vegetation (via fuel types). For instance, shrub fuel type enhance fire severity whereas adult mature conifer decrease the potential for high intensity fire.

We calibrated the fire regime using three attributes: fire rotation period, fire size distribution, and fire severity - (Keeley *et al.*, 2011). We calculated fire attributes through LANDSAT imagery and products derived from the Monitoring Trends in Burned Severity program (Eidenshink *et al.*, 2007). We chose the period 2000-2010 excluding the 2002 Biscuit Fire (ca. 202,000ha), because we wanted to discount the effect of one outlier fire that had a large influence over the fire regime (Fig S4 and Table S11). However, the targeted fire rotation period still portray the 1984-2010 period including such big fire. Fire severity was calculated as percent crown damage, and was approximated by using the relationship built in the study area between percent crown damage and the difference Normalized Burned Ratio Index (Thompson *et al.*,2007). Recent patterns of fires size, severity and frequency were reproduced by the model (Table S12; input parameters in Table S13 and S14).

We accounted for spatial differences in fire regimes by dividing the study area into three regions reflecting differences in contemporary ignition rates and fuel moisture (Fig S5 and Fig S6). These regions were identified by combining remote sensing estimates of the fog belt (Torregrosa *et al.*, 2016) and density estimators of fire occurrence as a distance from roads and human settlements (Figure S5). Fuel classifications were species and age specific and were derived from similar Mediterranean ecosystems studies (Syphard *et al.*, 2011).

## RESULTS

### Climate change scenarios and forest development under climate change

The climate change scenarios predicted consistent warming throughout the seasons ranging from 1.18 to 2.9 °C on average (Fig. 2a), whereas seasonal precipitation patterns varied between scenarios, GCMs, and seasons (Fig. 2b). Precipitation generally increased during the winter, with increases between 8 and 60 mm/month depending on the climate change scenario. Spring and fall had the highest temperature increase (rcp 8.5; Fig. 2). Conversely, summers were predicted to be hotter – ranging from 1.2 to 4.9 °C/month – with slightly lower precipitation – between 0 and 10 mm/month, depending on the scenario. These changes became more evident towards the latter half of the 21^st^ century simulation period.

**Figure 2:**
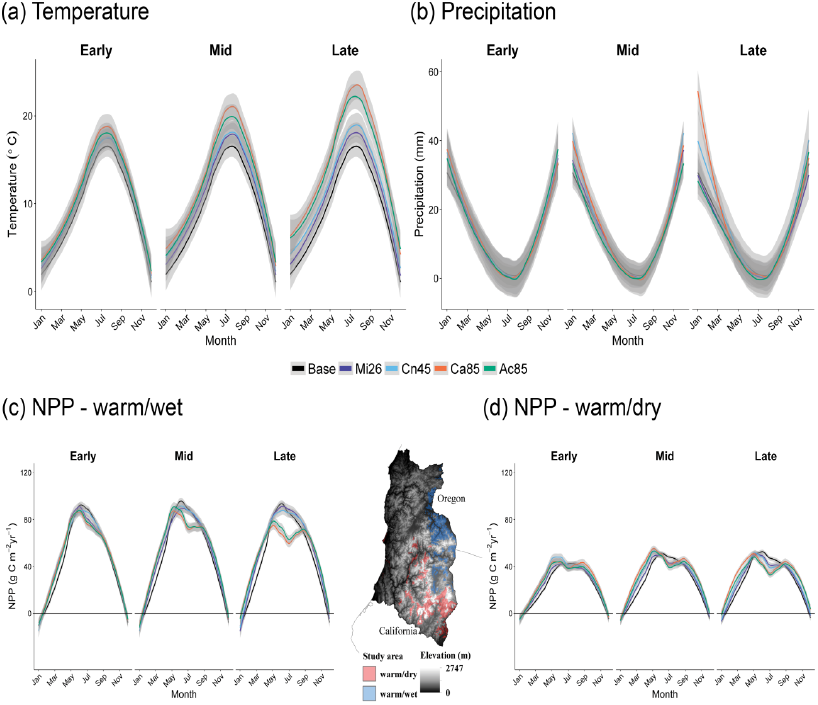
Climate seasonal regimes and the simulated effects on forest productivity. Temperature (a) and precipitation (b) under baseline and climate change conditions for the whole landscape. Effects of climate on forest net primary productivity (NPP) in two locations chosen to highlight the model’s response to environmental gradient under all climate scenarios:(c) warm-wet and (d) warm-dry. See Table 1 for climate change scenario acronyms.

Modeled forest growth rates reflected changes in soil water availability, which varied spatially depending on precipitation, temperature and soil characteristics (Fig. 2c and 2d). We selected two ecoregions that bound the gradient of water balance conditions in the area and thus show different responses to environmental conditions. In these ecoregions, modeled forest productivity ranged between 0.11 gC m^-2^yr^-1^ and 104 gC m^-2^yr^-1^. The model projected shifts in growth, with strong growth limitation during summer under climate change (Fig 2c and 2d). The model also projected a lengthening of the growing season, with greater growth enhancement at the beginning of the growing season (late winter/ early spring) than at the end (Fig 2c and d). It is important to note that these growth behavior – earlier growth in winter-spring and reduced growth in summer – across climate change scenarios are most apparent later in the 21^st^ century (Fig 2).

**Table 1.** Baseline and climate change scenario projections. Relative projections represent a qualitative analysis of Fig S1 for the purpose of synthesis. These labels are based on average annual statistics over the course of the simulation (85 years) for mean annual temperature and annual precipitation. Annual probability of establishment shifts across species in conifer and shrubland-chaparral-hardwood (SCH) species under different climatic conditions. These values represent averages across species and time. Percentages in brackets indicate the percentage of probability of establishment loss with respect to baseline conditions.

| Climate Scenario | Emissions scenario (RCP) | Climate model | Relative projections * | Fire Rotation Period *<sup>1</sup> | Annual establishment probability |  |
| --- | --- | --- | --- | --- | --- | --- |
|  |  |  |  |  | Conifers | SCH |
| Baseline |  |  |  | 104-123 | 0.23 | 0.3 |
| Mi26 | 2.6 | MIROC5 | Mild hot – wetter | 98-117 | 0.17 (26 %) | 0.28 (7 %) |
| Cn45 | 4.5 | CNRM-CM5 | Hotter – wetter | 91-116 | 0.18 (22 %) | 0.29 (3 %) |
| Ac85 | 8.5 | ACCESS | Much hotter - drier | 82-100 | 0.15 (35 %) | 0.25 (17 %) |
| Ca85 | 8.5 | CanESM2 | Much hotter - wetter | 82-99 | 0.14 (39 %) | 0.24 (20%) |

\*See Fig. S1 for quantitative outputs.
\*1 Range indicates 25<sup>th</sup> -75<sup>th</sup> percentile

The model projected a decline in the probability of establishment under climate change (Table 1) due to increased summer drought. The decline ranged from 26-39% depending on the climate change scenario for the conifer species and between 7-20% for the SCH group (Table 1). In general, probability of establishment was lower for the higher temperature forcing scenarios (Ac85 and Ca85 vs. Mi26 and Cn45). The probability of establishment was higher in Mi26 vs Cn45, which we attribute to the compensating factor of water in this case (water availability is higher in Cn45 than in Mi26; Table 1 and Fig. S1).

### Increased fire activity

Climate change enhanced fire activity (Figure 3). Fire rotation periods (FRP) were lowered by approximately 20 years on average for the highest forcing climate change scenarios (i.e. Ca85 and Ac85 Fig. 3a), whereas there was little change in the FRP under milder forcing scenarios (Cn4.5 and Mi2.6). There was high variability among simulations within each climate change scenario, especially for those climate change scenarios with the least forcing. Simulations suggest that, while averaged FRP may not be statistically different from baseline conditions in 100 years, the result was much higher risk of conditions conducing to higher fire conditions (fire weather indices) that can lead to a large shift in FRP.

**Figure 3:**
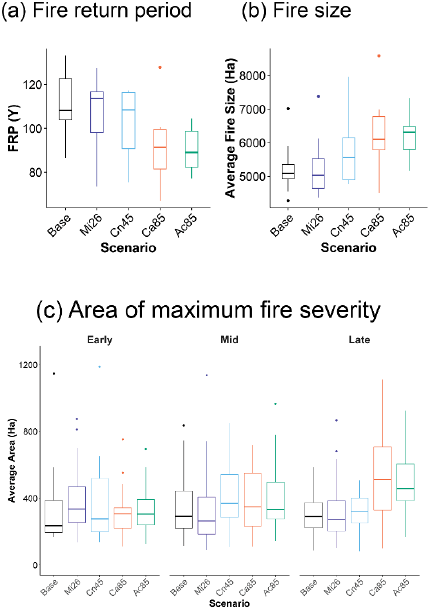
Fire regime model outputs: (a) Fire return period – time to burn an area of the same size of the area of study; (b) Fire Size for different simulation repetitions under baseline and climate change scenarios; (c) High fire severity area for different classes. Boxplot represent different the distribution of values across 9 simulation repetitions. See Table 1 for climate change scenario acronyms.

Average fire sizes increased under climate change (Fig. 3b), particularly under the warmest scenarios (Ca85 and Ac85). The model projected a maximum average increase of 1,011 ha and 1,222 ha between baseline conditions and Ca85 and Ac85 scenario, respectively. More importantly, climate change was projected to increase the severity of fires (Fig. 3c), especially in the last part of the century.

The number of megafires – above 100,000 ha (Fig. 4). Baseline conditions predicted the occurrence of large-scale fires of approximately 200,000 ha, while under the warmest climate change scenarios (Ac85 and Ca85) fires reached between 200,000 ha and 500,000 ha in several years. Only in the highest forcing scenarios did a fire reached a size beyond the half million hectares (1 out of 9 repetitions per scenarios in Ca85 and Ac85).

**Figure 4:**
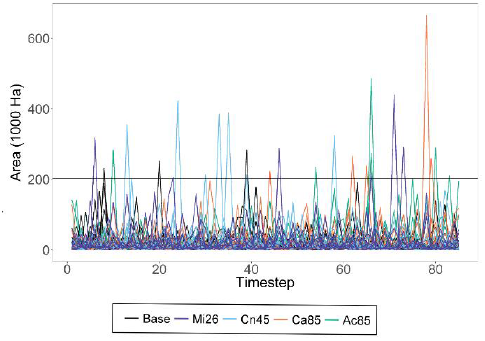
Time series of maximum fire size for different model simulation. Horizontal solid line indicates the historical maximum fire size recorded in the area (Biscuit fire 202,000 ha). See Table 1 for climate change scenario acronyms.

Mean fire return intervals maps highlight the spatial variability of fire recurrence (Fig. 5). Fire was more prevalent in the eastern side of the study area, consistent with the west to east gradient of increasing water deficit (Fig. 1a). The total area with mean fire return intervals under 25 years only increased between 1% and 5% of the study area depending on the climate change scenario (Fig. 5)

**Figure 5:**
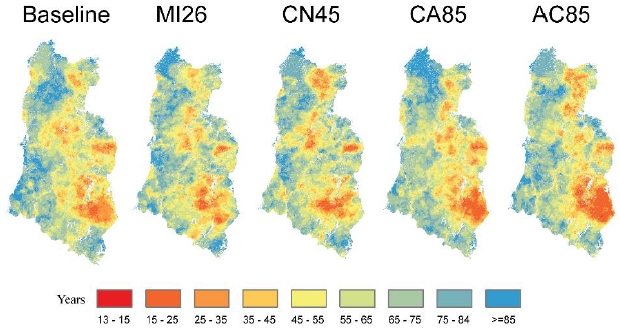
Spatial distribution of mean fire return intervals (MFRI) in the area. MFRI above 85 indicates that no fire was recorded in the area for the simulations analyzed. See Table 1 for climate change scenario acronyms.

### Dominance shifts

Compared to the current vegetation dominance pattern, simulations of both baseline and climate change conditions resulted in large-scale shifts in vegetation dominance, particularly within the drier eastern half of the study area (Fig. 6a). Major shifts were thus located where the model projected the highest fire activity. The most pronounced spatial disagreement between baseline and climate change simulations was concentrated in the center of the study area, coinciding with higher elevations and topographic complexity (Fig. 6b).

**Figure 6:**
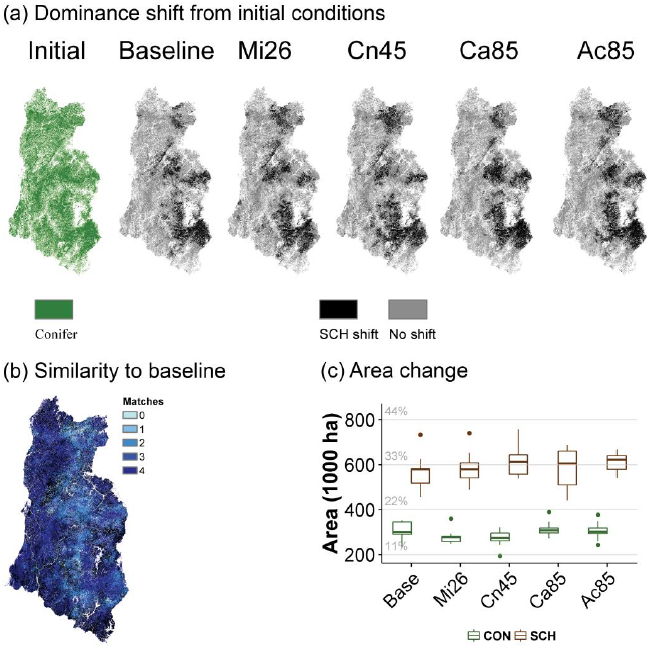
Shifts in forest type as determined by species dominance: (a) Forest dominance shifts compared to initial conditions; (b) Similarity index between climate change scenarios and baseline conditions. The index informs how many climate change scenarios agree with baseline (e.g. matches, 4 = maximum agreement, 0 = maximum disagreement); and (c) Area of conifer forest transitions remaining as conifer (CON) or shifted to shrubland-hardwood (SCH). See Table 1 for climate change scenario acronyms.

The model estimated a conifer dominance loss of more than half million hectares in areas affected by fire during 85 years of simulation, even under current (e.g. baseline; dominance loss 579,000 ha) conditions. That approximately represents 31% of the initial forest dominated by conifers in the study region. Climate change increased the area of conifer forest lost by an average of 7,000 ha when forced by the Mi26 scenario and 60,000 ha with the Ac85 scenario (Fig. 6c). Once more, the stochasticity of wildfire is highlighted in climate change scenarios that present higher precipitation under climate change (Fig 6c, height of boxplot in Cn45 and Ca85).

## DISCUSSION

Our results demonstrate a large potential for rapid vegetation shifts in the Klamath region during the 21^st^ century. Some of these shifts are driven by climate change, but simulations with current climate conditions also produced major shifts. Climate change, in the 100 year interval, may cause a decrease in the fire rotation period in the area and an increase in fire size.

These results concur with empirical and modeling studies that suggest that disturbances can rapidly accelerate vegetation shifts during climate change (De Frenne *et al.*, 2013; Serra-Diaz *et al.*, 2015; Stevens *et al.*, 2015) and suggest that rapid vegetation dominance shifts are likely to occur when strong feedbacks between co-existing species are present in the community (e.g. conifer forests and shrublands), as in the case of the Klamath with vegetation-fire interactions create alternative stable states. In our case, this synergy is reflected in the increase in fire severity along the simulation time, especially at the end of the 21^st^ century, coinciding with the expansion of shrubs and hardwoods (Fig 3c). Such potential for rapid shifts are likely to be found at biome transitions, such as the conifer forests and Mediterranean hardwoods of the Klamath, where several functional types and traits adapted to fire regimes coexist and climate conditions can lead to different vegetation community states. Our work suggests that the local empirical studies by Odion *et al.* (2010) and Tepley *et al.* (2017) can be generalized to a regional context, and portray a regional trend of increasing vulnerability of conifer forests to altered fire regimes along a west to east gradient with increasing aridity (Fig. 1a and Fig. 4b).

### Shifts in the 21^st^ century even under current climate conditions

One of the most striking results of the modeling experiment is that, on average, current baseline conditions (1949-2010) largely lead to the loss of almost 1/3 of the initial conifer forest extent.Several mechanisms could explain such results, but we argue that these stem from the combination of (i) disequilibrium of vegetation with disturbance dynamics.

Disequilibrium with climate refers to the lag between changes in abiotic conditions and the response of communities or species to such changes. Several lines of evidence suggest that species distributions and communities reflect past environmental changes rather than current environments (Sprugel, 1991; Donato *et al.*, 2012; Hughes *et al.*, 2013; Svenning & Sandel, 2013; Ordonez & Svenning, 2017). This emphasizes the need to stress the temporal scope of 100 years climate change projections. For instance, in a climate change model experiment, García-Valdés *et al.* (2013) found that forest species in Spain are predicted to increase their range and abundance under current climate as a result of last glacial maximum dynamics that pushed many species southwards in Europe. Specifically, in their study climate change would only affect that inertia in 3 out of 10 species. In our case, we cannot rule out the idea of the influence of the fire shortage during the Little Ice Age may have increased the dominance of forests of our region via fire shortage (Colombaroli & Gavin, 2010); and that legacy may be playing out today in a remarkably warmer climate after 1949-2010 (our climate baseline climate input). Such longer term dynamics may indeed obscure the short-term predictions of the consequences of climate change in forests.

In addition, the potential effects of recent historical fire suppression by humans, which has promoted conifer dominance in many forests (Parsons, & DeBenedetti, 1979; McIntyre *et al.*, 2015) may have exacerbated positive fire–vegetation feedbacks in certain landscapes, thereby facilitating extensive vegetation transformation (McWethy *et al.*, 2013; Tepley *et al.*, 2016). Indeed, empirical data indicate that such afforestation process may increase subsequent fire risk due to fuel continuity and exposure to high severity fires (Foule 2003, Miller 2009, Westerling 2016), at least at early stages of conifer development before negative forest feedbacks with fire arise. These un-intended effects together with rapid increases in temperature have led to an increase in fire sizes and the number of megafires in many regions (Adams, 2013), as simulated here in our study.

### Climate change signal at the end of the century: mechanisms acting in concert for rapid shifts coupled with long-term transitions

The interaction of several processes included in our modeling experiment indicated a low potential for maintaining the current level of conifer dominance in forests in the region, concurring with other mixed conifer forests in western United States that showed or predict major vegetation shifts (McIntyre *et al.*, 2015; Harvey *et al.*, 2016). Despite similarity to baseline conditions for half of the simulation our study, several projections suggest potential for longer term transitions. First, fire severity and fire sizes were projected to increase under climate change. Second, species probability of establishment decreased, particularly for conifer species; and third, the model suggests growth limitation, hampering post-fire regeneration.

The interaction of fire size and fire severity and fire return interval are key to understand post-disturbance and recolonization, and may have a strong bearing on the capacity of forest to recolonize burned patches (Harvey *et al.*, 2016; Johnstone *et al.*, 2016; Tepley *et al.*, 2017). Our simulations projected increased fire size, concurring with empirical work (Westerling *et al.*, 2006; Westerling, 2016). Particularly interesting is that the model suggested potential for an increase in megafires under climate change, surpassing the historical maximum fire sizes recorded in the area (Biscuit fire 202,000 ha). Similar projections based on statistical correlations also project an increase of megafires in North California (Barbero *et al.*, 2015). Interestingly, both fire size and severity seem to start differentiating from baseline at the end of the 21^st^ century, and the ‘slightly wetter’ scenario (Ca 85) produced higher fire activity in terms of size and severity (Fig.3) than the ‘slightly dryer’ scenario (Ac85). We can’t rule out the possibility that his effect may be due to the relatively low number of replications, but we argue that this is likely the effect of increased fuel build up due to wetter conditions in concert with a much warmer summer. Indeed, this may represent an early signal that under much dryer conditions negative feedbacks of vegetation with fire may arise in the future.

In addition to potential limitation of conifer re-colonization via dispersal, establishment probabilities were projected to decrease under climate change. This is reflected in the model through lower soil moisture and higher temperatures. Such influences appear to have played out after recent fires in the Klamath Mountains. Following high-severity fire, the trend of decreasing density of regenerating conifers with reductions in seed-source availability becomes steeper as climatic water deficit increases (Tepley et al. 2017). Thus, if climate change drives increases in the patch sizes for high-severity fire while also creating a more arid post-fire environment, dry portions of the landscape could face a substantial lengthening of the time to forest recovery after high-severity fire, increasing the probability that the post-fire SCH vegetation is perpetuated by repeated fire before it has a chance to return to forest cover (Coppoletta *et al.*, 2016). Similar decreases in opportunities for seedling establishment under drier conditions have been observed in western US forests (Davis *et al.*, 2016; Welch *et al.*, 2016), especially following high severity fires (Savage & Mast, 2005; Feddema *et al.*, 2013).Interannual climatic variation, which was large during the 21^st^ century, in the future may provide opportunities for species establishment even if average climatic conditions are not conducive to seedling establishment (Serra-Diaz *et al.*, 2016). However, high inter-annual climatic variation may also come at the cost of cohort mortality due to extreme dry conditions. On balance, the rate at which conifers successfully establish and persist for a decade or more in the Klamath declines with increasing aridity, with seed source proximity as a key interactive variable.

The model does not simulate drought-related cohort mortality due to processes of hydraulic failure, carbon starvation or a combination of both. It is thus likely that our results represent a conservative estimate of vegetation shifts. First, data shows an ongoing recurrence of tree mortality events in forests affected by drought (Allen *et al.*, 2010; Carnicer *et al.*, 2011), including the recent (2012-2015) major drought in the region (Asner *et al.*, 2016), and projections show that the probability of such droughts will increase in this region in the future. In addition, empirical data shows that larger trees may be more vulnerable to drought related mortality (Bennett *et al.*, 2015), which could further reinforce the transitions via fuel dryness enhancement (e.g. less shading) and increased seedling mortality due to less canopy shading.

Our model predicted an overall reduction in annual tree cohort growth, although there is high variation depending on the species, climate, and soils. Overall, and particularly for the drier regions in the landscape, summer growth is predicted to decline in both our model and across a network of tree-ring chronologies (Restaino *et al.*, 2016). But there is potential for additional growth in spring and to a lesser extent in the fall (Fig. 2b; see also Creutzburg *et al.*, 2016b).Indeed, earlier springs and overall extended growing seasons are important and could increase growth and thus the ability of conifers to recover after a fire (see Chmura *et al.*, 2011 and references therein) especially for the higher elevations in our study region. Using a combined measurements of flux tower and satellite information, Wolf *et al.* (2016) found that earlier spring could reduce the impact of a subsequent summer drought in 2012. However, the model outputs suggest that this effect may be transient and compensation may not hold during summer. Our results regarding growth should be interpreted with caution. Our model assumes full phenological adaptation, and it is likely that growth can be further constrained by maladaptations to the new phenological cycle (Morin *et al.*, 2009). The model also does not incorporate the potential effects of CO_2_ fertilization (Keenan *et al.*, 2011, 2013) that could speed up forest development and growth as well as alter successional dynamics (Anderson-Teixeira *et al.*, 2013; Miller *et al.*, 2016). The effects of CO_2_ on growth in this forest are subject to further scrutiny, and potential growth is likely to be limited by the low N deposition in the region (0.12 Kg N/ha annual average, NADP; <u>http://nadp.sws.uiuc.edu/</u>).

Our various model components highlight that different processes may hamper current conifer dominance. Indeed, the fire regimes that emerge from the interaction of vegetation, climate and soils show the high potential for developing long-term shrubland states (Lindenmayer *et al.*, 2011): Mean fire rotation intervals and higher fire severity projected in some areas are certainly too short (<= 40 years) to fully develop a mature conifer forest, concurring with the hypothesis of a general interval squeeze that may hamper the development of conifers (Enright *et al.*, 2015). Accordingly, empirical data show that broadleaf trees and shrubs typically comprise the majority of aboveground woody biomass for at least the first three decades following high-severity fire (Tepley et al. 2017), and when stands reburn before conifers regain dominance, fire-severity tends to be high (Thompson and Spies 2010, Lauvaux et al. 2016). Thus, successive high-severity fires on the order of decades could potentially preclude the recovery of conifer forests almost indefinitely.

### Average projections vs. Risk in disequilibrium systems

The differences in potential for conifer decline between simulations driven by baseline climatic conditions and those driven by climate change scenarios were, on average, somewhat less than might be anticipated (Fig. 4). Nevertheless, we argue that main differences between climate change and baseline conditions are better interpreted using the potential for extreme scenarios of shift rather than average conditions.

Some of the processes that have been analyzed may be of limited relevance in short time frames. For instance, lower probabilities of establishment in conifers will have a large repercussion on mature conifer forest abundance, but it is likely that they play a major role beyond the 21^st^ century given that forest development times operate at larger time scales. Also, the length of our simulations (85 years) comprises only about one complete fire rotation under each of the future climate scenarios. If high-severity fire is the mechanism that converts conifer forests to SCH, and repeated severe fire (possibly facilitated by positive fire–vegetation feedbacks) is the mechanism that perpetuates SCH once the conversion occurs, we may need a longer timeframe before we see these transitions play out extensively across the landscape. In a single fire rotation, some portions of the landscape burn more than once and others remain unburned, but the fire–vegetation feedbacks would have to be very strong to see large areas of conifer forest that are converted to SCH and then perpetuated in that state by repeated burning within a single fire rotation. Individual fire events are fast processes that can transform forests rapidly and may have major impacts at finer spatial scales and shorter temporal frames. However, the degree to which these effects persist may depend on slower processes of vegetation regrowth and the timing and severity of subsequent fires. Therefore, at this temporal span (~100 years, potentially one generation of trees), projections of vegetation shifts are more useful when interpreting the risk of such fires’ transformations rather than by averaged outputs.

Our modeling experiment projects a high potential for the loss of nearly one-third of the existing mature conifer forest across the Klamath region in the coming century, warning that there is potential for fast transitions in forests. Baseline model projections of widespread forest loss (Fig. 4 a,c) suggest that current widespread distribution of conifer forests are not in equilibrium with the late-20^th^ or early 21^st^-century conditions. Given this disequilibrium, the projected fire dynamics, and the increasingly challenging conditions for conifer to regenerate during climate change, this study highlights that current conifer forests may be holding a considerable amount of resilience debt (*sensu* Johnstone et al. 2016), likely to be paid during the 21^st^ century. Further research still needs to unveil to what extent local forest management aimed at reducing the vulnerability of conifer forests to severe fire, or facilitating their post-fire recovery could buffer against our projected forest loss, and to better understand how and to what degree we need to learn to adapt to the changing landscapes and disturbance regimes where preventing such changes may no longer be possible.

## ACKNOWLEDGEMENTS

This work was funded by the National Science Foundation program DEB-1353301. JMSD acknowledges further support from Microsoft Azure grant under the Climate Data Initiative. We thank Luca Morreale for technical help.

## SUPPORTING INFORMATION

Figure S1 Climate change scenarios considered.

Figure S2 Species specific biomass calibration

Figure S3 Plot level accuracy assessment

Figure S4 Total burned area for the period 1984-2010.

Figure S5 Cumulative ratio of fire ignitions from distance to road

Figure S6 Fire regions encompassing spatial variability in ignitions and spread rates.

Figure S7 Mean fire rotation thresholds on predictions of conifer dominance in forests.

Table S1 LANDIS-II general species parameters.

Table S2 Available light biomass table.

Table S3. Light establishment table.

Table S4. Century succession species parameters.

Table S5. Century succession functional group parameters.

Table S6. Initial ecoregion parameters.

Table S7. Ecoregion parameter table.

Table S8. Monthly maximum above-ground net primary productivity (ANPP) table (g m-2).

Table S9. Maximum biomass table.

Table S10 Fire statistics according to different time periods

Table S11 Fire regime calibration: Fire Size distributions and Severity.

Table S12 Shape parameters for fire size distributions for all climate models by fire region.

Table S13 Fuel types description and parameters for the Dynamic Fire and Fuel extension of

LANDIS-II.

